# A Cys2/His2 zinc finger protein acts as a repressor of gibberellins biosynthesis by regulating SD1/OsGA20ox2 in rice (Oryza sativa L.)

**DOI:** 10.1101/616433

**Authors:** Min Duan, Xiao-Juan Ke, Hong-Xia Lan, Xi Yuan, Peng Huang, En-Shun Xu, Xiu-Ying Gao, Ru-Qin Wang, Hai-Juan Tang, Hong-Sheng Zhang, Ji Huang

## Abstract

Gibberellins (GAs) play important roles in the regulation of plant growth and development. The green evolution gene *SD1* encoding gibberellin 20-oxidase 2 (GA20ox2) has been widely used in modern rice breeding. However, the molecular mechanism of how *SD1*/*OsGA20ox2* expression is regulated remains unclear. Here we report a Cys2/His2 zinc finger protein ZFP207 acting as a transcriptional repressor of *OsGA20ox2*. *ZFP207* was mainly accumulated in young tissues and more specifically in culm nodes. *ZFP207*-overexpression (*ZFP207OE*) plants displayed semi-dwarfism phenotype and small grains by modulating cell length. RNA interference of *ZFP207* caused higher plant and longer grains. The endogenous bioactive GA levels were significantly reduced in *ZFP207OE* plants and application of exogenous GA_3_ rescued the semi-dwarf phenotype. The *in vivo* and *in vitro* studies showed that ZFP207 repressed the expression of *OsGA20ox2* via binding to its promoter region. Together, ZFP207 acts as a transcriptional repressor of gibberellins biosynthesis and it may play a critical role in plant growth and development through fine-tuning GA biosynthesis in rice.

## Introduction

Plants adjust their growth or shapes under developmental and environmental signals and the plant hormones are the signal molecules during these processes. Gibberellins (GAs) are a class of phytohormones that modulate many aspects of plant growth and development, including seed germination, leaf expansion, stem elongation and flowering (Fleet and Sun, 2005). The genes encoding GA-biosynthetic enzymes have been identified in numerous species and are most highly expressed in rapidly growing tissues or organs (Olszewski *et al*., 2002). Only a small number of them, such as GA_1_ and GA_4_, are thought to function as bioactive hormones. Therefore, many non-bioactive GAs exist in plants as precursors for the bioactive forms or deactivated metabolites (Yamaguchi, 2008).

The synthesis of bioactive GAs is generally divided into two parts. The early steps contain that trans-geranylgeranyl diphosphate (GGDP) is converted and modified into GA_12_ by CDP synthase (CPS), ent-laurene synthase (KS), entkaurene oxidase (KO) and ent-kaurenoic acid oxidase (KAO) in plasmid (Sakamoto *et al*., 2004). The final stage of bioactive GA synthesis, from GA_53_/GA_12_ to GA_1_/GA_4_, is catalyzed through two parallel pathways (i.e. early-13-hydroxylation and non-13-hydroxylation pathways) by GA20-oxidase (GA20ox) and GA3-oxidase (GA3ox), two soluble 2-oxoglutarate-dependent dioxygenases (2ODDs) in the cytosol. Based on the mutant and expression analyses, the enzymes catalyzing the early steps in the GA biosynthetic pathway (i.e. CPS, KS, KO and KAO) are mainly encoded by single genes, while those for later steps (i.e. GA20ox, GA3ox, and GA2ox) are encoded by gene families. The major bioactive GAs are GA_1_, GA_3_, GA_4_ and GA_7_, while other GAs are mostly non-bioactive but the precursors for bioactive forms in plants (Yamaguchi, 2008).

Many components of GA signaling pathway have been identified in plants (Davière and Achard, 2013). The GA signal is perceived by the GA receptor, GID1, which forms the GA-GID1-DELLA complex. In absence of GAs, DELLAs accumulate and repress GA signaling while GAs promote the 26s proteasome-mediated destabilization of DELLAs, thereby the growth-inhibition by DELLAs are released (Dill et al., 2004). Modulation of GA biosynthesis or signaling pathway has a significant influence on plant growth and development. In rice, loss-of-function mutation of *GA20ox2* results in dwarf or semi-dwarf plants. The rice *semidwarf-1* gene is well known as the “green revolution gene.” This gene has contributed to the significant increase in crop production seen in the 1960s and 1970s, especially in Asia (Futsuhara and Kikuchi, 1997; Asano *et al*., 2011). Also, originally derived from the Chinese cultivar Dee-Geo-Woo-Gen (DGWG, also called Di-jiao-wu-jian), *SD1* provides rice cultivars with short, thick culms, improves lodging resistance and responsiveness to nitrogen fertilizer, resulting high yields without affecting panicle and grain quality (Monna *et al*., 2002). *SD1* is expressed in rapidly growing regions, such as leaf primordia, young leaves, elongating internodes, developing anthers, and embryo (Kaneko *et al*., 2003). This expression is also detected in reproductive meristems and stamen primordia during floral organogenesis. The expression pattern suggests that the enzymes encoded by rice *GA20ox2* play a major role in both vegetative and reproductive development.

Proteins containing the classical Cys_2_/His_2_ zinc finger domains are most abundant in eukaryotic genomes. Many of these proteins are transcription factors that function by recognition of specific DNA sequence (John *et al*., 2001). Proteins containing zinc finger domain(s) were found to play important roles in eukaryotic cells regulating different signal transduction pathways and biological processes, such as development and programmed cell death (Yilmaza and Mittlera, 2008). In the past decades, after zinc finger protein SUPERMAN modulating the number of stamens was identified and characterized (Bowman *et al*., 1992; Sakai, 1995), several Cys2/His2-type zinc finger proteins that contain single zinc finger domain have been shown to involve the regulation of plant growth or phytohormone metabolism. *KNUCKLES*, like *SUPERMAN*, encodes a small protein containing a single Cys2/His2 zinc finger and functions as a transcriptional repressor (Payne *et al*., 2004). *GIS*, *GIS2* and *ZFP8*, which encode Cys2/His2 transcription factors, were functionally interchangeable activators of trichome production and the corresponding genes have specialized to play distinct roles in GA and cytokinin responses during *Arabidopsis* development (Gan *et al*., 2007). Later on, another Cys2/His2 zinc finger protein *ZFP5* was identified to play an important role in controlling trichome cell development through GA signaling (Zhou, *et al*., 2011). *STAMENLESS1* (*SL1*), very closely related to *JAG* gene, has been proved to regulate floral organ identity in rice (Xiao *et al*., 2009). Huang *et al*. (2009) screened a large ethyl methane sulfonate (EMS)-mutagenized M2 rice population and isolated a *dst* mutant line with enhancer salt and drought tolerance compared to wide type. *DST* gene encodes a Cys2/His2 type zinc finger transcription factor with transcriptional active activity, which negatively regulates stomatal closure under salt and drought stress in rice by targeting to genes related to H_2_O_2_ homeostasis via binding to a *cis*-element sequence named DBS (DST binding sequence). Besides the role in stress response, DST also plays roles in the regulation of reproductive SAM activity in rice through controlling the expression of *Gn1a*/*OsCKX2* and enhances grain production (Li *et al*., 2013).

We have previously characterized a Cys2/His2 zinc finger protein ZFP207 in rice (Meng *et al*., 2010). ZFP207 contains a single Cys2/His2 zinc finger domain at N-terminal and a predicted EAR (ethylene associated region) motif (Hiratsu *et al*., 2002) at C-terminal. In this work, we find that ZFP207 acts as a transcriptional repressor of GA biosynthesis by targeting the green evolution gene *SD1*/*OsGA20ox2*. Our work provides a mechanism for the molecular regulations of *SD1*/*OsGA20ox2* expression and GA biosynthesis in rice.

## Materials and methods

### Plant material and growth conditions

Rice cultivar Zhonghua 11 (*Oryza sativa* L. subsp. *japonica*, ZH11) was used for all the experiments. Rice seeds were placed for 1 wk at 42°C to break any possible dormancy, soaked in the water at room temperature and then germinated for 1 d at 37°C. The most uniformly germinated seeds were sown in a 96-well plate from which the bottom was removed. The seedlings were cultured with Yoshida’s culture solution in the growth chamber with a light cycle of 14 h light at 28°C and 10 h dark at 23°C photoperiod. The seedling and root at the three-leaf stage, leaf blade, leaf sheath stem, node, and immature panicle at the heading stage from ZH11 were collected and immediately frozen in liquid nitrogen for further gene expression assay.

### Quantitative real-time PCR

The total RNA was isolated using the Trizol reagent (Invitrogen, USA) according to the manufacturer’s protocol. The first strand cDNA was synthesized with 2 μg of purified total RNA using the reverse transcription system (Promega, USA) according to the manufacturer’s protocol. Quantitative real-time PCR was performed on an ABI7500 using the SYBR Green method. The *18S rRNA* gene was used as an internal control (Jain *et al*., 2006). The data were analyzed according to the ΔΔCt method (Livak *et al*., 2001). Each analysis was performed with three replicates. The primers used for quantitative real-time PCR are listed in Table S1.

### Vector construction and rice transformation

For *ZFP207*-overexpression construct, the coding sequence of *ZFP207* was cloned into pCAMBIA1304 after the CaMV 35S promoter. For *ZFP207RNAi* construct, a *ZFP207* 204bp fragment apart from the conserved region was cloned into pTCK303 under the control of ubiquitin promoter. The BLAST program against NR database at GenBank (http://www.ncbi.nlm.nih.gov) was employed to analyze the specificity of RNAi. For *ZFP207-GFP* construct, the sequence of *ZFP207* was cloned into pCAMBIA1300221-GFP.1 between CaMV 35S promoter and GFP protein. For analysis of *ZFP207* expression, about 1.8 kb length of *ZFP207* promoter was amplified from rice genomic DNA and cloned into pCAMBIA1304 vector to create a fusion of the promoter and *GUS* (β-glucuronidase) reporter gene (*P_ZFP207_*::*GUS*). All resulted constructs were transformed into ZH11 via an *Agrobacterium*-mediated method as described before (Ge *et al*., 2004). Finally, three independent transgenic lines overexpressing *ZFP207* and two independent lines in which the expression of *ZFP207* was restricted were harvested, respectively. The generated *P*_ZFP207_::GUS transgenic plant was used to analyze the spatial-temporal expression of *ZFP207*. Primers used in vector construction are listed in Table S1.

### Trait measurements

The *ZFP207*-overexpression lines (OE1, OE13), *ZFP207*-RNA interference lines (R10, R17) and wide type ZH11 were chosen for each trait measurement. Until four-leaf period (about 25-d-old seedlings), 42 healthy individual plants of each line were picked and transferred to field environment until adult period. The grain traits and plant height were measured when the plants were completely matured under field environment. For plant height, panicle length and internode length, 15 individual plants of each line were selected and measured. After all the grains were harvested in the form of single plant, grains were dried at 42°C overnight and 10 random grains of each line were selected for grain length measurement.

### Histochemical analysis

Histochemical analysis was performed using the X-gluc buffer method (Couteaudier *et al*., 1993) with some modification. Different organs of *P_ZFP207_*::*GUS* transgenic seedling were incubated in a solution containing 50 mM NaP buffer at pH 7.0, 5 mM K_3_ Fe(CN)_6_, 5 mM K_4_ Fe(CN)_6_, 0.1% Triton X-100, and 1 mM X-Gluc, and incubated at 37°C for at least 12h. Primers used for *P_ZFP207_*::*GUS* construction are listed in Table S1.

### GA induction in second leaf sheath elongation

The effect of GA_3_ on second leaf sheath elongation was quantitatively determined as described previously (Matsukura *et al*., 1998) with some modifications. Seeds of *ZFP207OE* rice and wide type were surface-sterilized for 30 min with 0.3% NaClO solution, washed 6 times with sterile distilled water, soaked in 5 μM paclobutrazol for 24 h to eliminate background and then in sterile distilled water for an additional 48 h after washing out the paclobutrazol. Seeds were placed on a 1% agar plate with or without GA_3_ and incubated at 30 °C under continuous light. After 6 days, the second leaf sheath length was measured (n=6).

### Subcellular localization

The coding sequence of *ZFP207* was amplified and cloned into plant transient expression vector pA7 (Zhang *et al*., 2012) under the control of CaMV35S promoter to construct ZFP207::GFP fusion protein. The 35S::GFP fusion protein was used as control. The constructs were transferred into rice protoplast cells by polyethylene glycol (Chen *et al*., 2006). GFP fluorescent signals were detected using confocal laser scanning microscopy (Nikon ECLIPSE 80i, Japan). Primers used for subcellular localization are listed in Table S1.

### Transcriptional Activity Assay

The transcriptional activity assay of ZFP207 was performed in yeast cells and rice protoplast, respectively. The whole *ZFP207* coding region and the deletion fragment were cloned into pGBKT7 vector fused with GAL4 DNA-Binding domain to create the pGBKT7-ZFP207 and pGBKT7-ZFP207N (1-181a.a.) constructs. pGBKT7-ZFP179 and pGBKT7 were used as positive and negative controls, respectively. All these constructs were transferred into yeast strain AH109. The transcriptional activity assay was performed following the protocol of the manufacturer (Stratagene, USA).

For the constructions in rice protoplast, we used the pGL3-Basic vector (Promega, USA) with minor modification. Two *cis*-elements, 5×GAL4 (Sugano *et al*., 2003) and 3×DRE (Liu *et al*., 1998), and a minimal TATA promoter were cloned before luciferase reporter gene. Based on the construction of pGBKT7-ZFP207, the DBD (GAL4-DNA binding domain)-ZFP207 or -ZFP207N was amplified and cloned into pA7 under the control of CaMV35S promoter. To enhance the background expression level, 35S::OsDREB1A effector plasmid was constructed, which can bind to DRE *cis*-element was constructed (Dubouzet *et al*., 2003). The renilla luciferase was used as internal control. All the plasmids were transferred into rice protoplast by polyethylene glycol and cultivated overnight for the expression of luciferase genes. The luciferase activity was detected via GLOMAX 20/20 Luminometer (Promega, USA). Primers used here are listed in Table S1.

### Transient expression assay

Approximate 1500 bp length of *SD1*/*OsGA20ox2* and *OsKS1* promoters were amplified from rice DNA genome, respectively, and cloned into pGL3-Basic vector before minimal TATA promoter and luciferase reporter gene to construct reporter plasmid. *ZFP207* coding sequence was cloned into pA7 vector under the control of CaMV35S promoter to make 35S::ZFP207 effector plasmid. The renilla luciferase was used as the internal control. The protoplast transformation and luciferase activity are performed as above section. Primers used for transient expression assay are listed in Table S1.

### Chromatin immunoprecipitation (ChIP)-qPCR assay

ChIP was performed as previously described (Shu *et al*., 2013). The 35S:*ZFP207-GFP* and wild-type ZH11 seedlings were harvested and cross-linked in 1% formaldehyde for 30 minutes and then neutralized by 0.125M Glycine. After washing with sterilized water, the seedlings were ground in liquid nitrogen, and the nuclei were isolated. Immunoprecipitations were performed with the anti-GFP antibody and protein G beads. Chromatin precipitated with normal mouse IgG antibody was used as negative control. The quantitative real-time PCR analysis was performed using specific primers corresponding to the promoter region of *SD1*/*OsGA20ox2* with three replicates. The *18S rRNA* gene was used as an internal control. Primers used in ChIP assay were listed in Table S1.

### Computational and database analysis

Protein sequence similarity searches were performed in the NCBI (www.ncbi.nlm.nih.gov/BLAST). The multiple sequence alignment diagram was generated in ClustalX and GenDoc software. Sequence data from this article can be found in the GenBank/EMBL database under the following accession numbers: ZFP207 (Os07g0593000), DST (GQ178286), GmZFP1 (AM265629), KNU (AY612608), RBE (AB107371), SUPERMAN (AT3G23130), AtZFP1 (AT1G80730), AtZFP2 (AT5G57520), AtZFP5 (L39648), AtZFP8 (L39651), AtZFP10 (AT2G37740), AtZFP11 (AT2G42410), PROG1 (FJ155665), OsSL1 (EU443151), AtGIS (AT3G58070), AtGIS2 (AT5G06650).

## Results

### Expression patterns of ZFP207

ZFP207 is a Cys2/His2-type zinc finger protein showing homologous to many other single zinc finger proteins especially at the zinc finger domain (see Supplementary Fig. S1 at *JXB* online). Quantitative real-time PCR assay showed that *ZFP207* was more specifically expressed in rice stem, root and node compared to other tissues (Fig. 1a). To investigate the *in situ* localization of *ZFP207* mRNA, transgenic rice plants with a β-glucuronidase (GUS) construct driven by the *ZFP207* promoter was generated. The significant GUS signals were observed in multiple young tissues, such as coleorhiza, young culm and immature panicle. Especially, GUS signals were found with high levels at the culm nodes (Fig. 1b), implying the potential role for ZFP207 in the modulation of the internode elongation. The mRNA *in situ* hybridization assay revealed the strong signals of *ZFP207* at inflorescence meristems, especially stamen primordium and lateral root primordium (Fig. 1c). Taken together, *ZFP207* mainly locates at young tissues or those with active cell division, indicating that ZFP207 may involve the regulation of cell division or expansion during rice growth and development.

**Fig. 1.**
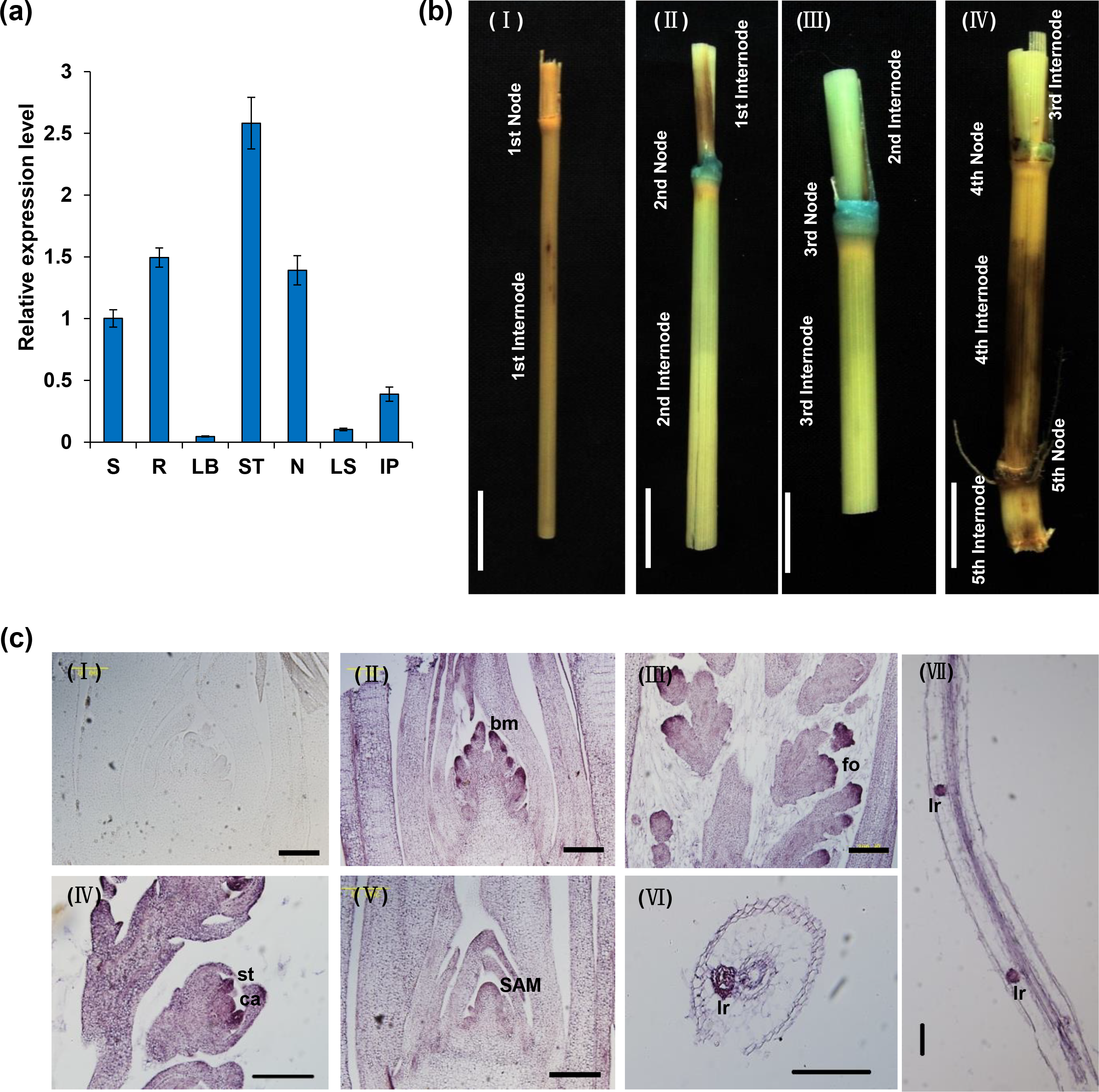
*ZFP207* expression pattern. (a) Quantitative real-time PCR analysis of *ZFP207* in various rice tissues. S, seedling; R, root; LB, leaf blade; ST, stem; N, node; LS, leaf sheath; IP, immature panicle. Data are presented as mean ± SE (n=3). (b) Histochemical *GUS* staining of the nodes of *ZFP207pro::GUS* transgenic rice internodes at mature stage, including the first internode and node site (I); the second internode and node site (II); the third internode and node site (III); the fourth and fifth internodes and node sites (IV). Bar=1 cm. (c) mRNA in situ hybridization of *ZFP207*. I, Spikelet meristem with sense probe. II, Spikelet meristem. III, Inflorescence meristem. IV, Floret meristem. V, SAM, stem axis meristem. VI, Transverse section of primary root. VII, Longitudinal section of primary root. Bar=100 μm. bm, branch meristem; fo, floral organ primordium; st, stamen; ca, carpel; lr, lateral root primordium.

### ZFP207 regulates plant height and internode elongation

To study the biological functions of *ZFP207* in rice, *ZFP207*-overexpression (*ZFP207OE*) and RNA-interference (*ZFP207RNAi*) transgenic plants were generated with a recipient *japonica* parent ZH11 (see Supplementary Fig. S2a at *JXB* online). We investigated the phenotypes of *ZFP207*-transgenic plants. It was found that *ZFP207OE* plants reduced the plant height at both seedling and adult stages and the panicle size compared to wild type ZH11 (Fig. 2a-e). By contrast, *ZFP207RNAi* rice exhibited the opposite phenotypes compared to *ZFP207OE* plants (Fig. 2a-e). As the rice height is mainly decided by the length and number of internodes, we found that the number of internodes was the same between ZH11 and *ZFP207*-transgenic plants. However, the internodes of *ZFP207OE* plants were shorter than ZH11, and those of *ZFP207RNAi* plants are increased in length especially for the first three internodes (Fig. 2f), indicating the critical role for ZFP207 in the modulation of the internode elongation. Scanning electron microscopy revealed that the cell length was reduced in *ZFP207OE* plants while increased in *ZFP207RNAi* plants (Fig. 2g-h), indicating the potential role for *ZFP207* in the regulation of cell elongation.

**Fig. 2.**
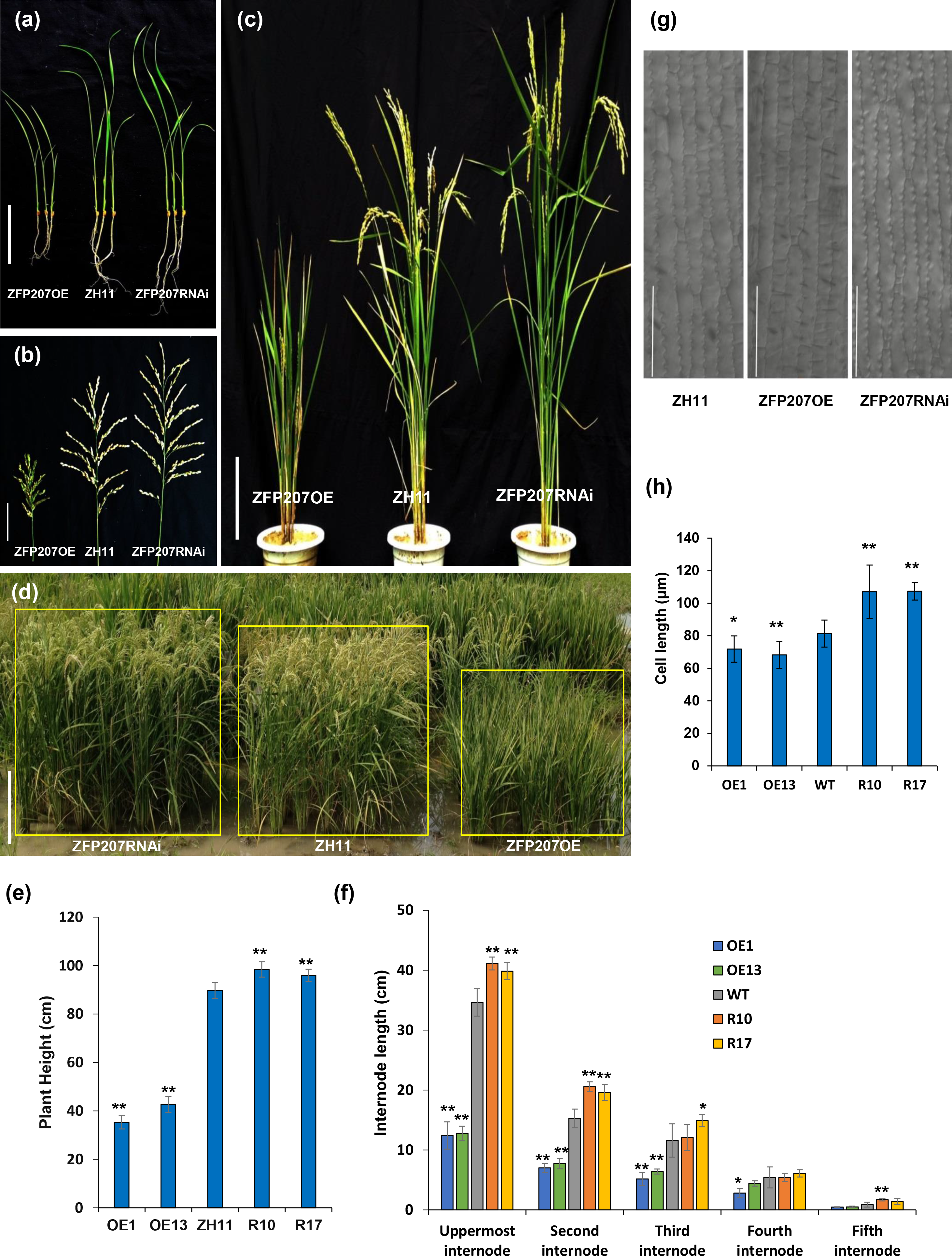
ZFP207 regulates plant height and internode elongation (a) The 15-d seedlings of *ZFP207OE* (S1), ZH11 and *ZFP207RNAi* (R10). Bar=5 cm. (b) Panicles of *ZFP207OE* (S1), ZH11 and *ZFP207RNAi* (R10) during mature period. Bar=5 cm. (c) Adult plants of *ZFP207OE* (S1), ZH11 and *ZFP207RNAi* (R10).Bar=20 cm. (d) Adult plants of *ZFP207OE* (S1), ZH11 and *ZFP207RNAi* (R10) grown in the field. Each yellow block showed that the specific line contained 42 individual rice plants. Bar=30cm. (e) Measurement of the plant height at mature period. (f) The length of internodes of *ZFP207OE* plants were shorter than ZH11, whereas *ZFP207RNAi* displayed longer panicle and internodes compared with ZH11. Bar= 5 cm. “*” and “**” represent significant differences compared to ZH11 with 0.01< *p* value< 0.05 and *p* value< 0.01, respectively. (g, h) The leaf cell size was examined and measured by differential interference contrast microscope. Bar=100 μm. “*” and “**” represent significant differences compared to ZH11 with 0.01< *p* value< 0.05 and *p* value< 0.01, respectively.

To further confirm whether the semi-dwarf phenotype of *ZFP207OE* plants relied on the expression level of *ZFP207*, we selected five individual plants from *ZFP207OE* lines with different dwarf levels and analyzed the *ZFP207* expression by semi-quantitative RT-PCR. As shown in Fig. S2b (see Supplementary Fig. S2b at *JXB* online), the height of *ZFP207OE* plants was related to the level of *ZFP207* expression, suggesting that the semi-dwarf levels were dose-dependent on *ZFP207* expression.

### ZFP207 modulates grain length and pollen viability

The grain length of *ZFP207OE* plants were reduced so that the grains became smaller compared to ZH11 grains. On the contrary, the *ZFP207RNAi* grains turned to be larger at grain length (Fig. 3a-b). Scanning electron microscope analysis showed that glume outer surface cells became smaller and closer in *ZFP207OE* grains, longer in *ZFP207RNAi* grains than in ZH11 (Fig. 3c-d). It was likely that the reduced grains of *ZFP207OE* plants resulted from a decrease in cell length.

**Fig. 3.**
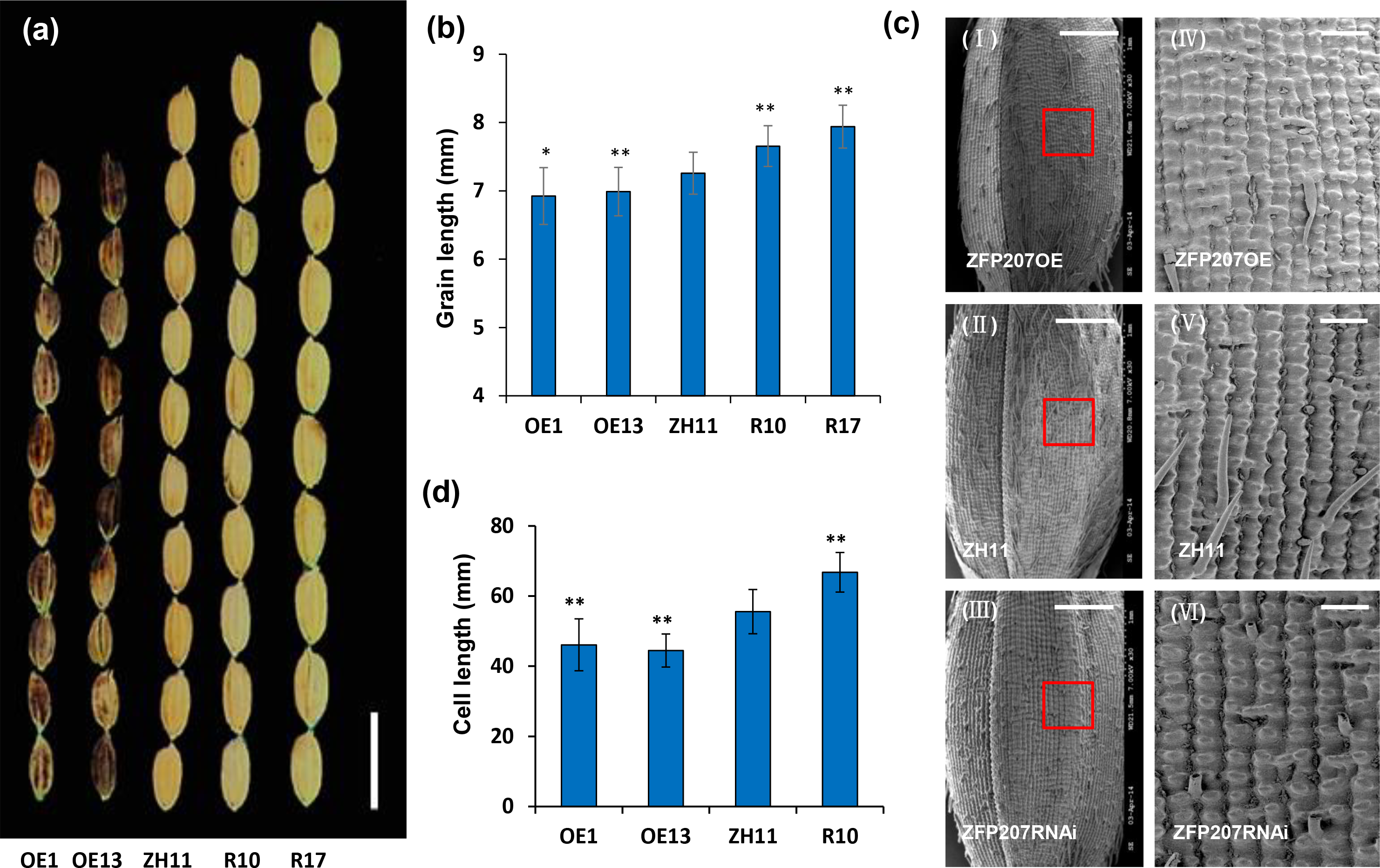
ZFP207 regulates grain length. (a) Mature seeds of *ZFP207OE*, ZH11 and *ZFP207RNAi* plants. Bar=1 cm. (b) Measurement of grain length for *ZFP207-*transnegic rice grains and wild type. (c) Scanning electron microscope (SEM) examination of seed epidermis of *ZFP207OE* (S1), ZH11 and *ZFP207RNAi* (R10) glume. (IV), (V), (VI) displays the amplification from the red boxes of (I), (II), (III) respectively. For (I), (II), (III), bar=1 mm. For (IV), (V), (VI), bar=100 μm. (d) Measurement of cell length of glume epidermis for *ZFP207* transgenic rice grains and wide type. “*” and “**” represent significant differences compared to ZH11 with 0.01< *p* value<0.05 and *p* value< 0.01, respectively.

As *ZFP207* is highly expressed in inflorescence meristems, *ZFP207* could influence the reproductive development. As expected, before flowering, *ZFP207OE* plants showed abnormal glumes and smaller stamens (see Supplementary Fig. S3a at *JXB* online). Meanwhile, we checked at least five visions under microscopy and found that the amount of pollen grains was significantly reduced in *ZFP207OE* plants and more than 50% pollen grains turned to be aborted (see Supplementary Fig. S3b-c at *JXB* online).

### ZFP207 negatively regulates GA biosynthesis

To determine whether the dwarf phenotype of *ZFP207OE* plants were due to GA deficiency or interrupted GA signaling, a GA rescue assay was performed. Our data showed that approximate 10^−5^ M exogenous GA_3_ could completely rescue the semi-dwarf phenotype of *ZFP207OE* seedlings, as indicated by the second leaf sheath length (Fig. 4a-b). In accordance with this result, we investigated endogenous GA levels in ZH11 and transgenic plants during three-leaf period, and found that bioactive GAs, such as GA_1_, GA_3_ and GA_4_ in *ZFP207OE* were significantly reduced compared to ZH11 and GA_3_, GA_4_ and GA_7_ showed the higher accumulations in ZFP207RNAi rice (Fig. 4c). These results indicate that ZFP207 represses the biosynthesis of bioactive GAs.

**Fig. 4.**
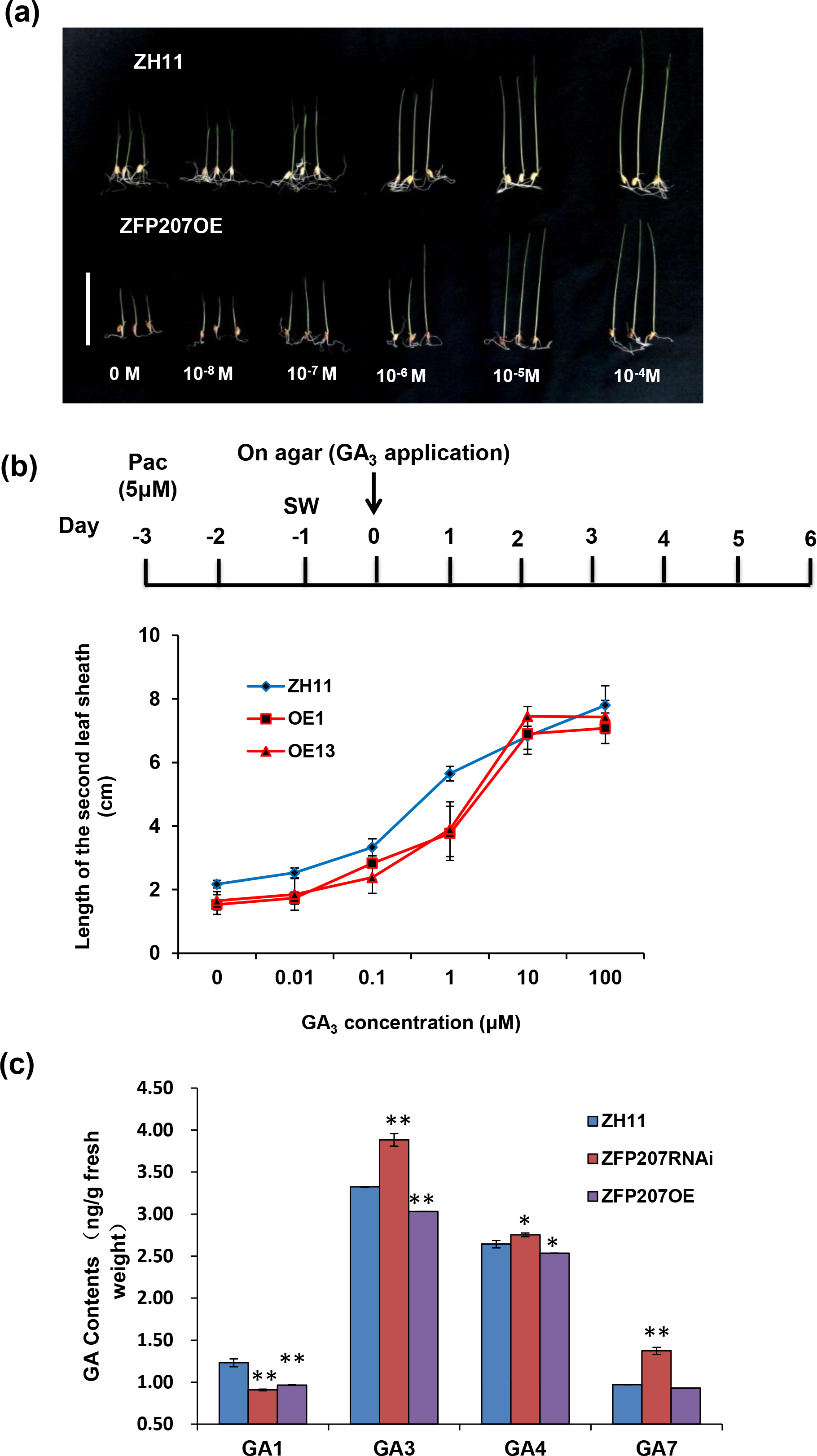
ZFP207 represses GA biosynthesis. (a, b) GA_3_ induced the second leaf sheath elongation. The second leaf sheath of *ZFP207OE* (S1) and ZH11 turned to be longer after the treatment with different concentrations of GA_3_. Bar=5 cm. PAC, Paclobutrazol; SW, seed washing. (c) Measurement of endogenous bioactive GAs in three-leaf period seedlings of *ZFP207OE* (S1), *ZFP207RNAi* (R10) and ZH11. Data are means ± SD from three trials. “*”,“**” represent significant differences compared with ZH11 with *p* value < 0.05 and *p* value < 0.01, respectively.

### ZFP207 negatively regulates expression of SD1/OsGA20ox2

Since ZFP207 is a Cys2/His2 zinc finger transcription factor, it is interesting to investigate whether ZFP207 regulate GA biosynthesis by targeting the genes encoding GA biosynthesis-related enzymes. We detected the expression of genes involved in GA synthesis and signaling in *ZFP207*-transgenic rice plants by quantitative real-time PCR. As shown in Fig. 5a, the expressions of GA synthesis genes, such as *OsKS1* and *SD1/OsGA20ox2* were markedly reduced in *ZFP207OE* lines as compared to ZH11 and increased significantly in *ZFP207RNAi* plants. Especially, the mRNA level of *SD1/OsGA20ox2* was highly decreased in *ZFP207OE* plants compared to in ZH11 and highly enhanced in *ZFP207RNAi* plants. On the other hand, expression of the genes related to GA signaling, such as the GA receptor *OsGID1*, *OsGID2* (Gomi *et al*., 2004; Ueguchi-Tanaka *et al*., 2005), and DELLA protein gene *OsSLR1* (Ikeda *et al*., 2001) were not markedly changed.

**Fig. 5.**
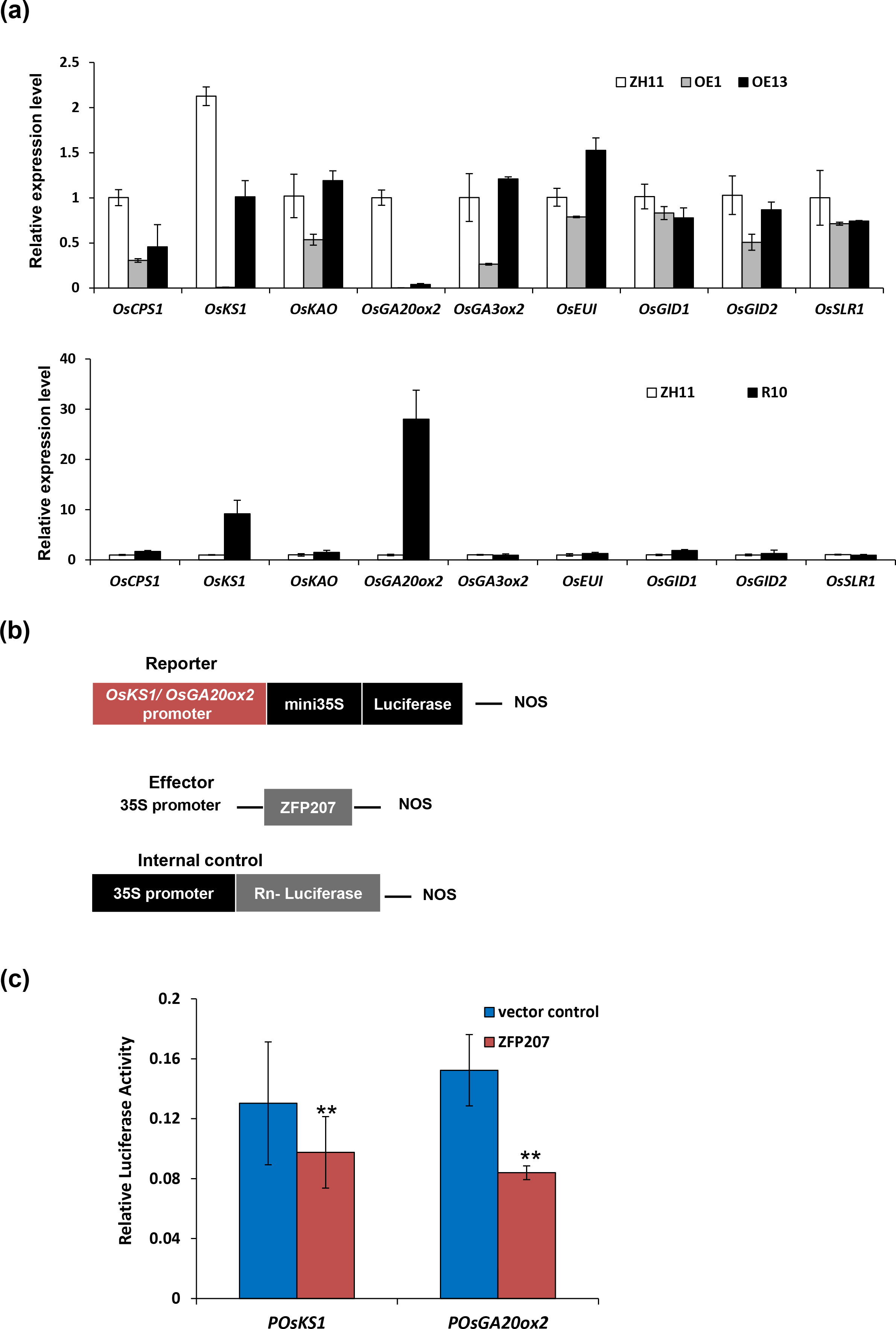
ZFP207 modulates expression of GA biosynthesis-related genes. (a) Quantitative real-time PCR analysis of expression of GA biosynthesis- and signaling-related genes in *ZFP207OE* (above) or *ZFP207RNAi* transgenic rice. (b) Construction of reporter, effector and internal control vectors for luciferase activity assay. (c) Measurement of luciferase activities. Vector control contains reporter and internal control plasmids; ZFP207 contains reporter, internal control and 35S::ZFP207 protein plasmids. “**” represents significant difference compared to vector control with *p* value<0.01.

Dual luciferase reporter assay system was used to reveal that whether ZFP207 could repress expressions of *SD1*/*OsGA20ox2* and *OsKS1 in vivo*. Protoplasts were transformed with effector plasmid and reporter plasmid containing promoter of *SD1*/*OsGA20ox2* or *OsKS1* and minimal cauliflower mosaic virus (CaMV) 35S promoter in upstream of a luciferase gene. The effector plasmid was composed of 35S promoter fused to the *ZFP207* coding sequence (Fig. 5b). Compared with vector control, luciferase activities were decreased after incubation with ZFP207 (Fig. 5c), confirming that ZFP207 performed the repression activity of the expression of *SD1*/*OsGA20ox2* and *OsKS1* in rice.

### ZFP207 acts as a transcriptional repressor

The subcellular location of ZFP207 was determined by transient expression in rice protoplast. The ZFP207-GFP fused protein was expressed in rice protoplast and fluorescent signal of ZFP207-GFP was specifically detected in nucleus, whereas the GFP control was distributed throughout the whole protoplast cell (Fig. 6a). Nuclei localization of ZFP207 was further confirmed by DAPI (4’, 6-diamidino-2-phenylindole)-staining.

**Fig. 6.**
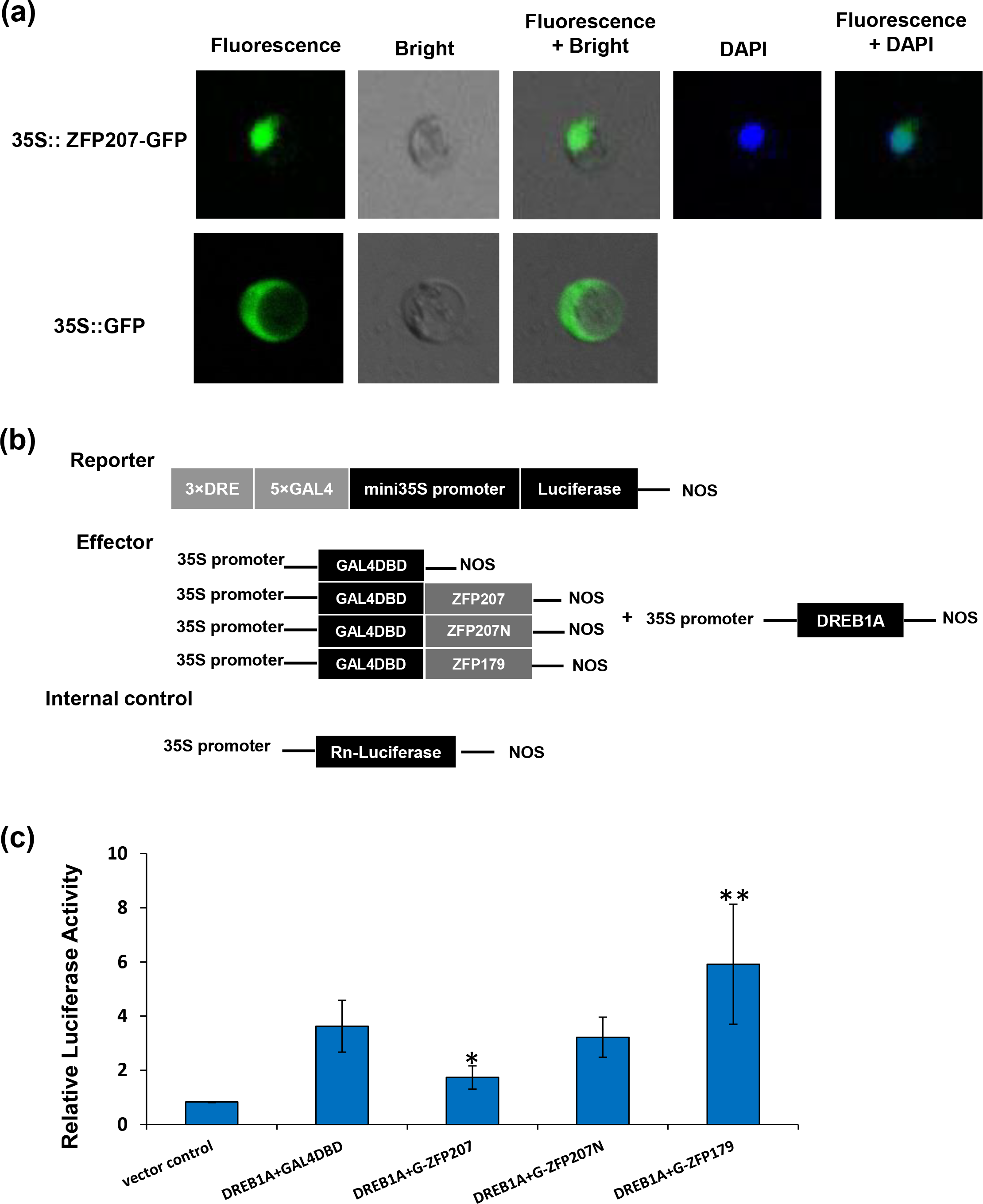
ZFP207 acts as a transcriptional repressor. (a) Subcellular localization of ZFP207-GFP fusion protein in rice protoplast cells. (b) Construction of reporter, effector and internal control vectors for luciferase activity assay. (c) Measurement of luciferase activity. Vector control contains reporter and internal control plasmids; DREB1A+GAL4DBD contains reporter, internal control, 35S::DREB1A protein and 35S::GAL4DBD protein plasmids; DREB1A+G-ZFP207 contains reporter, internal control, 35S::DREB1A protein and 35S::GAL4DBD-ZFP207 fusion protein plasmids; DREB1A+G- ZFP207N contains reporter, internal control, 35S:: DREB1A protein and 35S::GAL4DBD- ZFP207ΔEAR fusion protein plasmids; DREB1A+G-ZFP179 contains reporter, internal control, 35S::DREB1A protein and 35S::GAL4DBD-ZFP179 fusion protein plasmids. “*” represents significant difference between DREB1A+GAL4DBD and DREB1A+G-ZFP207, 0.01< *p* value < 0.05; “**” represents significant difference between vector control and DREB1A+G-ZFP179, *p* value < 0.01.

Yeast hybrid system was performed to investigate whether ZFP207 has transcriptional activation activity. Deletion analysis showed that not only the whole coding sequence of ZFP207, but ZFP207 partial sequences fused with GAL4 transcription factor binding domain (BD) couldn’t grow on SD/-Trp-His-Ade plate (see Supplementary Fig. S4 at *JXB* online), suggesting that ZFP207 has no transcriptional activation activity at least in yeast cells.

To further analyze whether ZFP207 had transcriptional repression activity, we used dual luciferase reporter assay system to examine the transcriptional activity of ZFP207 in rice protoplasts. Protoplasts were transformed with effector plasmids and reporter plasmid containing GAL4, DRE fragments and minimal cauliflower mosaic virus (CaMV) 35S promoter in upstream of a luciferase gene. Several effector plasmids were used in this assay (Fig. 6b). The GAL4DBD-ZFP179 (Sun *et al*., 2010) was used as the positive control and 35S::Rn-luciferase was used as internal control. As shown in Figure 6c, co-expression of DREB1A and GAL4DBD proteins in protoplasts showed induction in the expression of the luciferase reporter gene as compared to vector control. Compared with co-expression of GAL4DBD and DREB1A, co-expression of DREB1A and GAL4DBD-ZFP207 displayed significant down-regulation in the expression of the luciferase reporter gene. When GAL4DBD fused with ZFP207N, the repression activity of ZFP207 was fully abolished, suggesting that ZFP207 protein lacking C-terminus lost the transcriptional repression activity. Taken together, ZFP207 functions as a transcription repressor in rice cells.

### ZFP207 targets SD1/OsGA20ox2

As *SD1*/*OsGA20ox2* was dramatically decreased in *ZFP207OE* and increased in *ZFP207RNAi* plants, it would be interesting to investigate whether *SD1*/*OsGA20ox2* is directly regulated by ZFP207. Firstly, dual luciferase reporter assay system was used to reveal that whether ZFP207 could repress expression of *SD1*/*OsGA20ox2 in vivo*. Protoplasts were transformed with effector plasmid and reporter plasmid containing promoter of *SD1*/*OsGA20ox2* and minimal cauliflower mosaic virus (CaMV) 35S promoter in upstream of a luciferase gene. The effector plasmid was composed of 35S promoter fused to the *ZFP207* coding sequence (Figure 7a). Compared with vector control, luciferase activity was decreased after incubation with the ZFP207 expressed protein (Fig. 7b).

**Fig. 7.**
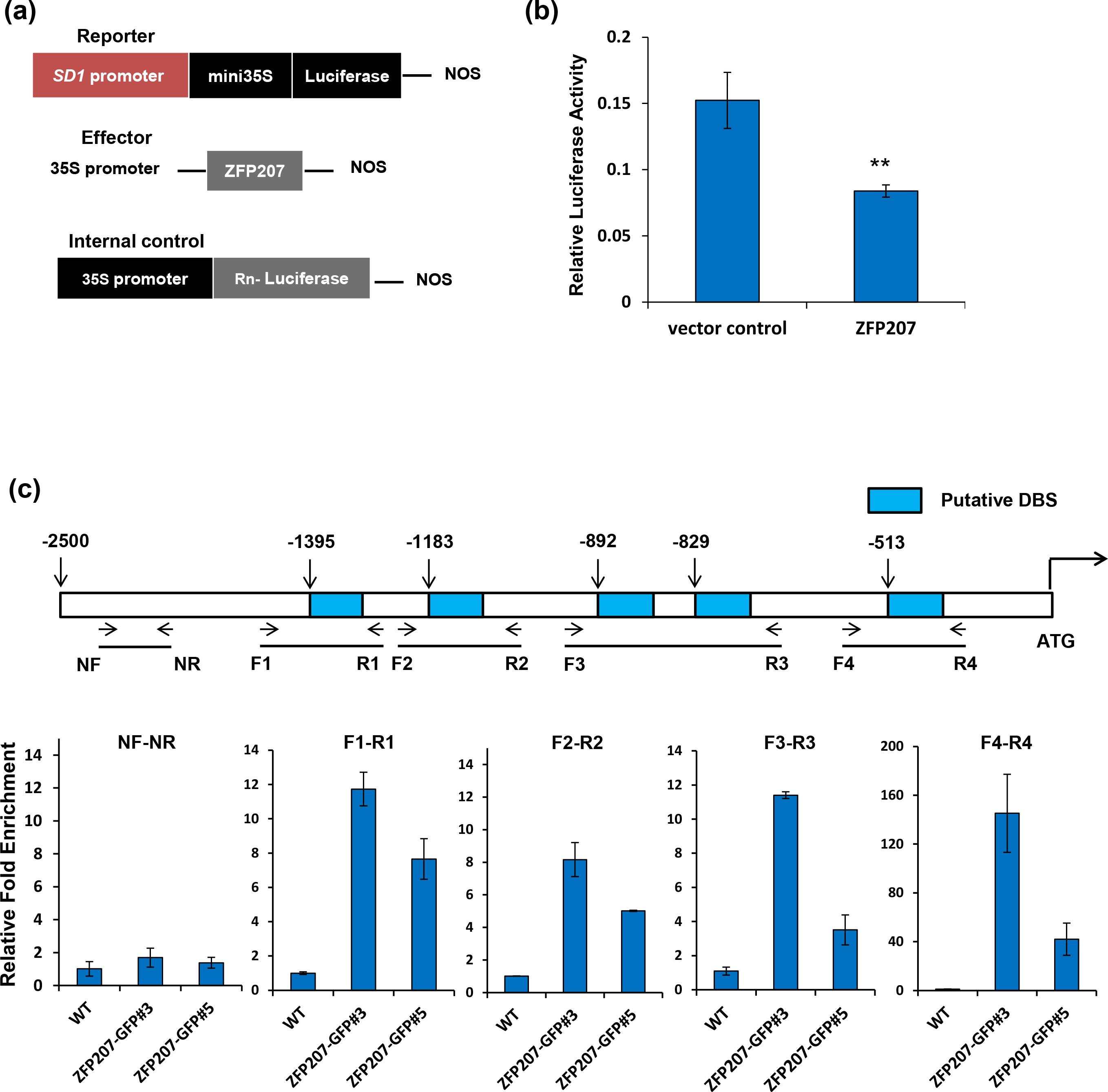
ZFP207 represses *SD1/OsGA20ox2* expression by targeting its promoter. (a, b) ZFP207 represses *SD1/OsGA20ox2* expression *in vivo*. (a) Construction of reporter, effector and internal control vectors for luciferase activity assay. (b) Measurement of luciferase activity. Vector control contains reporter and internal control plasmids; ZFP207 contains reporter, internal control and 35S::ZFP207 protein plasmids. “**” represents significant difference compared to vector control with *p* value<0.01. (c) ChIP-qPCR assay. Two 35S:ZFP207-GFP transgenic lines, *ZFP207-GFP*#3 and *ZFP207-GFP*#5 were generated and used in this assay. Primers used in the ChIP-qPCR assay are indicated by arrows and shown in Table S1. Total protein extracts from 35S:ZFP207-GFP transgenic plants grown for two weeks were immunoprecipitated with an anti-GFP antibody. Chromatin precipitated with normal mouse IgG antibody was used as internal control, and wild-type ZH11 was used as negative control. Data are means ± SD from three biological replicates.

Previous study has revealed that DST, a Cys2/His2 type zinc finger transcription factor in rice, could bind to a conversed TGNTANN(A/T)T sequence named DBS (DST binding sequence) (Huang *et al*., 2009). To verify the direct interaction of ZFP207 with *SD1/OsGA20ox2* promoter *in vivo*, we performed a ChIP (chromatin immunoprecipitation) assay with the *ZFP207-GFP* transgenic plants. As expected, the phenotype of 35S::*ZFP207-GFP* transgenic plants was similar with *ZFP207OE* plants (data was not shown). ZFP207-bound fragments enriched by immunoprecipitation with anti-GFP antibody were used for quantitative real-time PCR analysis. As shown in Fig. 7c, we chose four parts containing putative DBS regions and found that all these fragments were significantly enriched in two independent *ZFP207-GFP* transgenic lines. The highest enrichment was found at the F4/R4 region in the nearest place to the *ZFP207* translational start codon and no markedly enrichment was found at the region without DBS.

## Discussion

### ZFP207 is a GA biosynthesis repressor

Plant development is finely regulated and coordinated through the action of several classes of plant hormones, in which GAs are important to stimulate growth of most organs through enhanced cell division and cell elongation (Colebrook *et al*., 2014). Therefore, the precise modulation of GA content is critical to maintain the normal plant growth and organ development and less or more GA level usually results in the developmental disorders. The green revolution gene *SD1* encodes a GA 20-oxidase (OsGA20ox2), which regulates the synthesis of biologically active gibberellins (GAs), (Monna *et al*., 2002; Spielmeyer *et al*., 2002; Sasaki *et al*., 2002). Loss of function of *OsGA20ox2* caused semi-dwarfism phenotype. Obviously, *OsGA20ox2* gene plays a critical role in GA biosynthesis.

In the present study, we discovered a Cys2/His2-type zinc finger protein ZFP207 as a novel regulator of GA biosynthesis in rice by targeting *SD1*/*OsGA20ox2*. *ZFP207* is predominately expressed in the cell-active tissues, like culm nodes, where GAs are usually accumulated and cells are actively dividing and elongating. ZFP207 is therefore crucial to maintain the GA abundance at an adequate level that ensures the normal plant growth and development through modulating GA biosynthesis. Accordingly, with the increase of *ZFP207* expression, abundance of active GAs are significantly reduced and the GA-deficiency phenotypes were observed in *ZFP207OE* plants, including semi-dwarfism, reduced length of root and panicle, and reduced grain size as well as decreased pollen viability. Most of these phenotypes could be also observed in GA biosynthesis or signaling deficiency mutants (Chhun *et al*., 2007; Li *et al*., 2011; Ji *et al*., 2014). Similarly, a NAC transcription factor OsNAC2 was found to negatively regulate GA pathway by directly modulating the expression levels of *OsEATB* and *OsKO2* encoding the repressors of GA biosynthesis and signaling in rice (Chen et al., 2015). Although in *ZFP207OE* plants, the expression of *SD1*/*OsGA20ox2* was largely decreased, the phenotypes of *ZFP207OE* were not identical to *sd1* mutant, implying some other genes might be also directly or indirectly regulated by ZFP207.

### ZFP207 is involved in internode elongation and grain length

Plant height is an important agronomic trait for crops. Expression of *ZFP207* was highly accumulated in rice culm nodes, implying that ZFP207 might regulate internode elongation, which finally decides the plant height. Todaka *et al*. (2012) found that a phytochrome-interacting factor-like protein OsPIL1 highly expressed in nodes functioned as a key regulator of internode elongation, thereby affected the plant height. However, since OsPIL1 is not related to GA biosynthesis, the regulatory pathway of internode elongation may vary for ZFP207 and OsPIL1. More genetic and biochemical experiments are required to clarify whether there is a cross-talk of ZFP207 and OsPIL1. The GA biosynthesis and signaling has been described as necessary for internode elongation in rice (Ayano *et al*., 2014). The morphological and cellular comparisons among wild-type, *ZFP207OE* and *ZFP207RNAi* suggested that ZFP207 modulates cell expansion. Therefore, ZFP207 could modulate the internode elongation by restricting cell expansion. For ideal plant architecture, the adequate plant height confers not only lodging resistance but enough plant biomass. Thus the modulation of plant height is also considered in breeding practice. Besides ZFP207, several regulators including AP2/ERF transcription factors (Qi *et al*., 2011; Magome *et al*., 2004), A20/AN1 zinc-finger protein (Zhang *et al*., 2016) and bZIP proteins (Fukazawa *et al*., 2011) were involved in regulating plant height though modulating GA biosynthesis and/or signaling. These regulators modulate the GA signaling majorly through direct or indirect regulating the expression levels of genes responsible for GA biosynthesis or signaling.

Besides the impact on plant height, alteration of *ZFP207* expression resulted in some other phenotypic changes, including ferity and grain length. Unlike other factors such as *GS3* (Fan *et al*., 2006; Mao *et al*., 2010) and *OsPPKL1* (Zhang *et al*., 2012) that affect cell proliferation, our data showed that knock-down of *ZFP207* increased the seed size through modulating cell length. By contrast, *ZFP207OE* plants produced smaller cells and grains. Since *ZFP207* was highly expressed in the meristems of the reproductive organs, ZFP207 might function in regulating the reproductive organ size through modulating GA levels. It would be interesting to improve rice yield and/or appearance quality if the expression of *ZFP207* could be operated through genetic engineering approaches. However, even the grain size was increased in *ZFP207RNAi* plants, the grain weight were not significantly increased. DST (drought and salt tolerance) is a ZFP207-like zinc finger protein with transcriptional activation activity (Figure S1; Huang *et al*., 2009). Unlike the role of ZFP207 on GA regulation, DST regulates the phytohormone cytokinin (CK) level by modulating expression of *Gn1a/OsCKX2* which controls the grain number in rice (Li *et al*., 2013).

### ZFP207 directly regulates SD1/OsGA20ox2

*SD1*/*OsGA20ox2* is critical to GA biosynthesis and known as green evolution gene. However, the molecular mechanism of how *SD1*/*OsGA20ox2* expression is regulated remains unclear. *SD1*/*OsGA20ox2* expression was largely reduced in *ZFP207OE* plants but higher in *ZFP207RNAi* plants, suggesting that ZFP207 is an upstream repressor of *SD1*/*OsGA20ox2*, which is the first transcription factor reported so far as the modulator of *SD1*/*OsGA20ox2* expression. Our studies suggest that ZFP207 functions as a transcriptional repressor and directly targeting *SD1*/*OsGA20ox2* promoter region. However, there are many Cys2/His2-type zinc finger proteins showed transcriptional activation activities, like ZFP182 (Huang *et al*., 2012) and ZFP179 (Sun *et al*., 2010). DST, which contains a single zinc finger domain like ZFP207, is also a transcriptional activator (Huang *et al*., 2009). Unlike these zinc finger proteins, ZFP207 showed the transcriptional repression activity. Our data showed that ZFP207 binds to DST binding site (DBS)-like sequences which are present in the promoter region of *SD1*/*OsGA20ox2*, suggesting the direct control of ZFP207 on *SD1*/*OsGA20ox2* expression. Besides ZFP207, several transcription factors also participate in the regulation of GA biosynthesis in rice, like OsbZIP58 (Wu *et al*., 2014), GDD1 (Li *et al*., 2011), OsEATB (Qi *et al*., 2011) and OsNAC2 (Chen *et al*., 2015). Although TF-mediated regulatory and interaction network of GA biosynthesis is not clarified, these TFs coordinately regulate the GA biosynthesis through targeting various GA biosynthetic genes and modulate plant growth and development in rice.

## Acknowledgements

This work was supported by National Key Research and Development Program (2016YFD0100400), Natural Science Foundation of China (31571627), the Fundamental Research Funds for the Central Universities (KYZ201804), Jiangsu Collaborative Innovation Center for Modern Crop Production and Cyrus Tang Seed Innovation Center, Nanjing Agricultural University.

## Supplementary data

Supplementary data are available at JXB online.

**Supplementary Fig. S1** The comparison of amino acid sequences among ZFP207 and other plant zinc finger proteins.

**Supplementary Fig. S2** Expression analysis of *ZFP207* in transgenic rice.

(a) Quantitative real-time PCR analysis of expression of *ZFP207* in *ZFP207OE* and *ZFP207RNAi* transgenic rice.

(b) The dwarfism of *ZFP207OE* plant is related to *ZFP207* expression level. Bar=20cm.

**Supplementary Fig. S3** ZFP207 affects pollen viability.

(a) Florets of *ZFP207OE* (S1), ZH11 and *ZFP207RNAi* (R10) before flowering. White block in (IV), (V), (VI) showed that the anthers of *ZFP207OE* were smaller than ZH11 and *ZFP207RNAi*. Bar=1 cm.

(b, c) The pollen viability analysis of *ZFP207*-transgenic rice by potassium iodide staining. Bar=10 μm. Data in (c) are means ± SD from five visions.

**Supplementary Fig. S4** Transcriptional activity assay of ZFP207 in yeast cells.

ZF, zinc finger; EA, EAR-motif. ZFP179 (Sun *et al*., 2010) served as a positive control.

**Supplementary Table S1** Primers used in this study.

## Author contributions

J.H. planned and designed the research. M.D., X.K., H.L., X.Y., P.H., E.X., R.W., X.G. and H.T. performed the research. M.D., H.Z. and J.H. analyzed the data. M.D., H.Z. and J.H. wrote the article.

